# Cyclase-associated protein 2 (CAP2) controls MRTF-A localization and SRF activity in mouse embryonic fibroblasts

**DOI:** 10.1101/2020.10.19.344929

**Authors:** Lara-Jane Kepser, Laura Soto Hinojosa, Chiara Macchi, Massimiliano Ruscica, Elena Marcello, Carsten Culmsee, Robert Grosse, Marco B. Rust

## Abstract

Recent studies identified cyclase-associated proteins (CAPs) as important regulators of actin dynamics that control assembly and disassembly of actin filaments (F-actin). While these studies significantly advanced our knowledge of their molecular functions, the physiological relevance of CAPs largely remained elusive. Gene targeting in mice implicated CAP2 in heart physiology and skeletal muscle development. Heart defects in CAP2 mutant mice were associated with altered activity of serum response factor (SRF), a transcription factor involved in multiple biological processes including heart function, but also skeletal muscle development. By exploiting mouse embryonic fibroblasts (MEFs) from CAP2 mutant mice, we aimed at deciphering the CAP2-dependent mechanism relevant for SRF activity. Reporter assays and mRNA quantification by qPCR revealed reduced SRF-dependent gene expression in mutant MEFs. Reduced SRF activity was associated with moderately increased G-actin levels and reduced nuclear levels of MRTF-A, a transcriptional SRF coactivator that is shuttled out of the nucleus and, hence, inhibited upon G-actin binding. Moreover, pharmacological actin manipulation restored MRTF-A distribution in mutant MEFs. Our data are in line with a model in which CAP2 controls the MRTF-SRF pathway in an actin-dependent manner. While MRTF-A localization and SRF activity was impaired under basal conditions, serum stimulation induced nuclear MRTF-A translocation and SRF activity in mutant MEFs similar to controls. In summary, our data revealed that in MEFs CAP2 controls basal MRTF-A localization and SRF activity, while it was dispensable for serum-induced nuclear MRTF-A translocation and SRF stimulation.

## Introduction

Cyclase-associated proteins (CAPs) have been recognized as actin-binding proteins (ABP) two decades ago (Balcer et al. 2003; Bertling et al. 2004; Freeman and Field 2000; Hubberstey and Mottillo 2002), but significant progress in their molecular function has been achieved only recently (Chaudhry et al. 2010; Jansen et al. 2014; Johnston et al. 2015; Kotila et al. 2018; Kotila et al. 2019; Mu et al. 2019; Shekhar et al. 2019). These studies unraveled a role for yeast and mammalian CAPs in disassembly of actin filaments (F-actin) and in the ATP-for-ADP-exchange on actin monomers (G-actin) that is essential for F-actin assembly. Hence, CAPs emerged as important regulators of F-actin dynamics, the spatiotemporally controlled assembly and disassembly of F-actin. While these studies advanced our knowledge of their molecular functions, the physiological relevance of mammalian CAPs largely remained elusive, also because appropriate animal models were lacking. This holds true specifically for the ubiquitously expressed family member CAP1 (Jang et al. 2019), while recent studies revealed arrhythmia, cardiac conduction defects as well as dilated cardiomyopathy in systemic and heart-specific CAP2 mutant mice (Field et al. 2015; Peche et al. 2012; Stockigt et al. 2016). Additionally, skeletal muscle development and myofibril differentiation was retarded in systemic CAP2 mutants, which displayed a myopathy characterized by a large number of ring fibers associated with motor function deficits (Kepser et al. 2019). Together, these studies emphasized a pivotal role for CAP2 in striated muscles, in agreement with its abundant expression in heart and skeletal muscle (Field et al. 2015; Kepser et al. 2019; Peche et al. 2007). Heart defects in CAP2 mutant mice were associated with altered gene expression including an upregulation of genes whose expression is controlled by serum response factor (SRF), and they were partially restored upon SRF inhibition (Xiong et al. 2019), suggesting a causal relationship between SRF dysregulation and heart pathology. However, the mechanism underlying increased SRF activity upon CAP2 inactivation remained unknown.

SRF is a ubiquitously expressed and highly conserved transcription factor that was first identified in studies of fibroblast serum response. Its activity is mainly regulated by two classes of transcriptional coactivators, namely the ternary complex factors (TCFs) as well as myocardin and myocardin-related transcription factors (MRTFs) (Olson and Nordheim 2010).

TCFs and MRTFs compete for a common surface domain on the DNA-binding domain of SRF, interact with SRF in a mutually exclusive manner and activate different sets of SRF-target genes (Esnault et al. 2014; Gualdrini et al. 2016). The TCF-SRF pathway is promoted by Ras and mitogen-activated protein kinases (MAPKs), induces expression primarily of immediate early genes (IEG) and has been implicated in cell-cycle re-entry and proliferation (Gualdrini et al. 2016). Instead, the MRTF-SRF pathway is controlled by the availability of G-actin and induces expression of cytoskeleton-related genes (Esnault et al. 2014). Specifically, G-actin binds to MRTF and promotes its translocation into the cytosol, thereby inhibiting MRTF-SRF-dependent gene expression (Olson and Nordheim 2010). Hence, the actin regulator CAP2 may control SRF activity via a mechanism that involves actin and MRTF.

By exploiting mouse embryonic fibroblasts (MEFs) from systemic CAP2 mutant mice, we here i) tested whether CAP2 controls SRF activity in cell types other than heart cells and ii) aimed at deciphering the underlying mechanism. Reporter assays and quantitative PCR (qPCR) revealed reduced SRF activity in MEFs upon CAP2 inactivation. Impaired SRF activity in CAP2 mutant MEFs was associated with moderately increased G-actin and reduced nuclear MRTF-A levels. Further, MRTF-A distribution in mutant MEFs was normalized by pharmacological actin manipulation. While our data were in line with a role for CAP2 in regulating SRF activity via the actin-MRTF-A pathway in non-stimulated MEFs, serum stimulation equally induced nuclear MRTF-A translocation and SRF activity in control and CAP2 mutant MEFs.

## Material and Methods

### Preparation and culture of MEFs

MEFs were isolated from CAP2-KO mice and CTR littermates at E12.5 as previously described (Rehklau et al. 2012; Seiler et al. 2008). Briefly, dissected embryos were minced and treated with 0.25% trypsin (Invitrogen) after removal of the head and organs. After 15 min of incubation at 37°C, tissue was pulled through a 0.9-mm needle to obtain isolated cells. After washing twice with serum-free Dulbecco’s modified Eagle’s medium (DMEM; Invitrogen), isolated cells were cultured in DMEM containing 10% fetal calf serum (FCS; PAA). Primary MEFs (termed passage 0) were grown for 1 day. On the next day, MEF immortalization was started according to the 3-day transfer protocol (3T3) immortalization protocol that defined immortalization as completed after passage 12. MEFs were kept at 37°C and 5% CO_2_ in DMEM containing 4.5 g/l glucose and 10% FCS, as well as penicillin (100 m/ml; PAA) and streptomycin (100 mg/ml; PAA). MEFs were passaged every 2–3 days. All experiments were performed with MEFs of P12–P40.

### Generation of MEF cell line stably expressing MRTF-A-GFP

MRTF-A-GFP was stably expressed in MEF cells (control and CAP2 mutant cell lines). For that, first HEK293T cells were transfected using the calcium phosphate method. For lentivirus production, HEKT293T cells were cotransfected with the lentiviral packaging vectors psPAX and pMDG.2 together with the pInducer-MRTF-A GFP plasmid as described before (Hinojosa et al. 2017). After 48 h, supernatants containing viral particles were harvested, filtered, and used to transduce MEF cells. Transduced MEF cells were selected by FACS-based cell sorting. Expression of MRTF-A–GFP from pInducer20 was induced by 333 ng/ml doxycycline.

### Live cell imaging

To analyze MRTF-A subcellular localization upon serum stimulation, MEF cells stably expressing pIND20-MRTF-A-GFP were serum-deprived for 48 h and then live cell imaging was performed. Cells were stimulated 24 h before imaging with 333 ng/ml doxycycline to induce MRTF-A-GFP expression.

Serum stimulation was performed adding 20% serum directly to the cells under the microscope to follow the effects of the stimulation on MRTF-A translocation over time. Microscopic imaging was performed using confocal laser-scanning microscope (LSM 800, Carl Zeiss) and a 63x 1.4 NA oil objective lens (Carl Zeiss). Time-lapse microscopy was performed at 37°C in a CO_2_-humidified incubation chamber (Pecon, CO_2_, module S1) using ZEN software (Carl Zeiss).

### SRF-luciferase reporter gene assay

To assess SRF activity, we generated MEF cell lines (control and CAP2 mutant) expressing the firefly luciferase reporter where MRTF-SRF promoter 3Da.luc was linked to GFP. To generate the MEF cell lines stably expressing the MRTF-SRF luciferase reporter, first HEK293T cells were transfected using the calcium phosphate method. For lentivirus production, HEKT293T cells were cotransfected with the lentiviral packaging vectors psPAX and pMDG.2 together with the lentiviral vector FUGW expressing MRTF-SRF promoter 3Da.luc linked to GFP. Generation of the lentiviral luciferase reporter construct has been described before (Hinojosa et al. 2017). After 48 h, supernatants containing viral particles were harvested, filtered, and used to transduce MEF cells. Transduced MEF cells were selected by FACS-based cell sorting.

MEF cells expressing the MRTF-SRF luciferase reporter were serum-deprived overnight and stimulated either with or without 20% serum for 24 h or 48 h. Then, cells were lysed with 200 μl Triton lysis buffer on ice and collected in 1.5 ml Eppendorf tube, followed by 10 min centrifugation at 13 000 rpm at 4°C. The amount of firefly luciferase was measured luminometrically for each condition using a Glomax 96 Microplate Luminometer (Promega) (Baker and Boyce 2014).

### Nucleofection

To study MEF morphology and CAP2 localization, MEF cells were transfected with either 3μg pEGFP-C1 (Clontech) or 3μg pEGFP-C1-MmCAP2 (kindly provided by Elena Marcello) using the 4D Nucleofector (Lonza) with the Amaxa P3 Primary Cell 4D Nucleofector X Kit (Lonza) according to the manufacturer’s instructions.

### Fixation and Immunohistochemistry

MEFs were plated on coverslips coated with 0,01 % calf skin collagen (Sigma Aldrich) in 0,1 M acetic acid. Coverslips were fixed with 4% PFA in PBS for 10 min and afterwards washed three times with PBS for 5 min each.

For immunohistochemistry, coverslips were treated for 1 hour with blocking solution and afterwards incubated over night at 4°C in carrier solution containing the primary antibody. After three washing steps of 5 min in PBS, coverslips were incubated for 2 hours at room temperature (RT) in carrier solution containing Alexa-488 conjugated secondary antibodies (1:200, Life Technologies). For phalloidin and DNase I treatment, coverslips were incubated for 2 hours at RT with both conjugates in PBS. Afterwards, coverslips were counterstained for 10 min at RT with the intercalating dye Hoechst 33342 (Invitrogen) diluted 1:1,000 in PBS. After two washing steps in PBS, coverslips were fixed on microscopic slides with Aqua-Poly/Mount (Polyscience Inc.). Images were acquired with a Leica SP5 confocal microscope using 20x and 40x objectives (N.A. 0.7, N.A. 1.3, respectively). Images were processed with Fiji software (ImageJ 1.51w) and analyzed by exploiting the cell counter plug-in.

Primary antibodies used for immunohistochemistry: mouse anti-MRTF-A (G8, Santa Cruz). To label F- and G-actin, we exploited rhodamine (1:200, R415, Invitrogen) phalloidin and Alexa-488 DNAse I conjugate (1:500, D12371, Invitrogen).

### Treatment with LATB and JASP

Previous to treatment with LATB (Abcam, #ab1442091) or JASP (Abcam, #ab141409), pIND20-MRTF-A-GFP control and CAP2 mutant cells were activated 24 h before with 333 ng/ml doxycycline. Cells were treated for 4 h with either 25nM LATB, 25nM JASP or an equal volume of DMSO as control.

### Immunoblot analysis

Immunoblots were performed as described before (Kepser et al. 2019). Briefly, generation of total protein from MEF cells was contained by homogenization of cells in lysis buffer containing protease inhibitor (Complete, Roche). For determination of G/F-actin ratio fractionation was performed as described in Huang et al. 2013.

Protein extracts were denatured at 95°C in Laemmli buffer, separated by SDS-page and blotted onto a polyvinylidene difluoride membrane (Merck) by using a Wet/Tank Blotting System (Biorad). Membranes were blocked for 1 hour and afterwards incubated with primary antibodies in blocking solution over night at 4°C. As secondary antibodies, horseradish peroxidase (HRP)-conjugated antibodies (1:20,000, Thermo Fisher Scientific) were used and detected by chemiluminescence with ECL Plus Western Blot Detection System (GE Healthcare). Primary antibodies used for immunoblots: rabbit anti-CAP2 (1:1,000, #15865-1-AP, Proteintech), mouse anti-actin (1:5,000, #NB600-535, Novus Biologicals), mouse anti-GAPDH (1:10,000, #MAB5718, R&D Systems)

### Quantitative PCR

Total RNA from MEFs was isolated using peqGold Trifast (VWR) according to the manufacturer’s instructions. To exclude DNA contamination, samples were treated with the TURBO DNA-free kit (Invitrogen), followed by reverse transcription using iScript cDNA synthesis kit (Bio-Rad) according to the manufacturer’s protocol. Quantitative PCR (qPCR) was performed in the STEP-One Light cycler (ABI Systems) using the iTaq SYBR Green Supermix (Bio-Rad) for detection of target genes. Three technical replicates were averaged and normalized to GAPDH in order to determine mRNA levels. Relative changes were calculated using the ΔΔCt method. Primer sequences: SRF (forward (f): TTCCCGTCCGAGGAAACAT, reverse (r) GGCTCTTTTGACCCAGACCAT), Vinculin (f: AGCCCAGATGCTTCAGTCAGA, r: GGTCAGATGTGCCAGAAAGGA), c-Fos (f: TTCCTACTACCATTCCCCAGCC, r: GATCTGCGCAAAAGTCCTGTG, GAPDH (f: CCCTTCATTGACCTCAACTA, r: CCAAAGTTGTCATGGATGAC), ACTA2 (f: GGCATCCACGAAACCACCTAT, r: CTGTGATCTCCTTCTGCATCCT), EGR2 (f: TGCTAGCCCTTTCCGTTGA, r: TCTTTTCCGCTGTCCTCGAT), CYR61 (f: AATCGCAATTGGAAAAGGCA, r: TGAAAAGAACTCGCGGTTCG).

### Cell viability

Cell proliferation was evaluated by using electrical impedance monitoring, through the xCELLigence Real-Time Cell Analysis (RTCA; Roche Diagnostics) system. When cells proliferate, the current flow between the microelectrodes, placed in the bottom surface of each well of a 96-well plate, is impeded. The impedance of this electron flow is reported as arbitrary cell index-values. For this assay, CTR and KO MEF cells were seeded in a 96-well plate (6,000 cells/well) and treated or not with erastin (0.7μM; Calbiochem).

Metabolic activity as an indicator for cell viability was assessed through the 3-[4,5-dimethylthiazole-2-yl]-2,5-diphenyltetrazolium bromide (MTT) assay. In viable and metabolically active cells, MTT is reduced to a purple formazan. CTR and KO MEF cells have been seeded in 96-well plates (2,000 cells/well) and treated with erastin (0.7μM) for 8 hours. Subsequently, MTT (0.5mg/mL; Merck) was added for 1 hour at 37°C for the purple formazan production. Absorbance was measured at 570-630nm with FLUOstar (BMG Labtech).

### Mitochondrial morphology

CTR and KO MEF cells were seeded in 8-well ibidi slides (Ibidi GmbH) at a density of 7,000 cells per well, stained with MitoTracker Deep Red FM (200nM for 30 minutes at 37°C; Invitrogen) and fixed with 4% paraformaldehyde for 20 minutes at room temperature. Images were acquired using a Leica DM6000 epi-fluorescence microscope (63x objective), by using an excitation wavelength of 620 nm and detecting emission using a 670 nm filter (red). Mitochondrial shape was classified as described before (Grohm et al. 2010): (i) category I comprises cells with healthy, elongated and equally distributed mitochondria, which are organized in a tubular network; (ii) category II comprises cells with partially fragmented mitochondria, which are still distributed throughout the cytosol, (iii) category III comprises cells with completely fragmented mitochondria, accumulating around the nucleus. At least 500 cells were counted by an experimenter blinded to the genotype.

### Mitochondrial superoxide formation

CTR and KO MEF cells, seeded in 24-well plate (10,000 cells/well), have been treated or not with erastin (0.7μM) for 8 hours. Subsequently, stained with MitoSOX Red (1.25μM; Invitrogen) for 30 minutes at 37°C, cells were harvested for FACS analysis (excitation 488 nm, emission 690/50 nm; Guava easyCyte Flow Cytometer, Merck). Increased red fluorescence has been correlated with the formation of mitochondrial reactive oxygen species (ROS). Data were collected from at least 5,000 cells and four replicates per condition.

### Lipid peroxidation

CTR and KO MEF cells, seeded in 24-well plate (10,000 cells/well), have been treated or not with erastin (0.7μM) for 8 hours. Subsequently, stained with BODIPY 581/591 C11 (2μM; Thermo Fisher Scientific) for 1 hour at 37°C, cells were harvested for subsequent FACS analysis (excitation 488nm, emission 525/30nm and 585/50nm). The shift from red to green fluorescence was used to analyze lipid peroxidation. Data were collected from at least 5,000 cells and four replicates per condition.

### Mitochondrial membrane potential

Mitochondrial membrane potential was analyzed by using the MitoPT TMRE Kit (ImmunoChemistry Technologies). CTR and KO MEF cells were seeded in 24-well plate (10,000 cells/well), treated or not with erastin (0.7μM) for 16 hours. Cells were stained with TMRE (0.2μM) for 30 minutes at 37°C and harvested for subsequent FACS analysis (excitation 488nm, emission 690/50nm). A decrease of TMRE fluorescence was representative of a loss of mitochondrial membrane potential. Data were collected from at least 5,000 cells and four replicates per condition.

### Statistical analysis

Statistical analysis was performed using the Prism statistical analysis package (GraphPad Software). Data are expressed as mean ± standard error (SE) for all main figures and mean ± standard deviation (SD) for the supplementary figure. All experiments have been conducted in at least three independent experiments. For supplementary figure eight replicates per condition were used if not specified. Differences between groups were evaluated by either one- or two-way ANOVA followed by post-hoc Dunns’ or Scheffé’s test, Chi-square test or student’s t-test and considered significant at p < 0.05. All experiments were conducted by experimenters blind to the genotype.

## Results

### CAP2 inactivation moderately increases G-actin levels in mouse fibroblasts

To study the cellular function of CAP2 in mammalian cells, we generated immortalized mouse embryonic fibroblasts (MEFs) from two CAP2^−/−^ mice (termed KO) and two CAP2^+/+^ control littermates (CTR) at embryonic day (E) 12.5. Immunoblots confirmed absence of CAP2 from both KO MEF lines (Fig. 1A). Due to the lack of specific antibodies suitable for immunocytochemistry, we examined subcellular CAP2 localization in MEFs by expressing a green fluorescent protein (GFP)-tagged CAP2 construct. GFP-CAP2 was homogenously distributed within the cytosol and largely absent from the nucleus (Fig. 1B).

**Figure 1.**
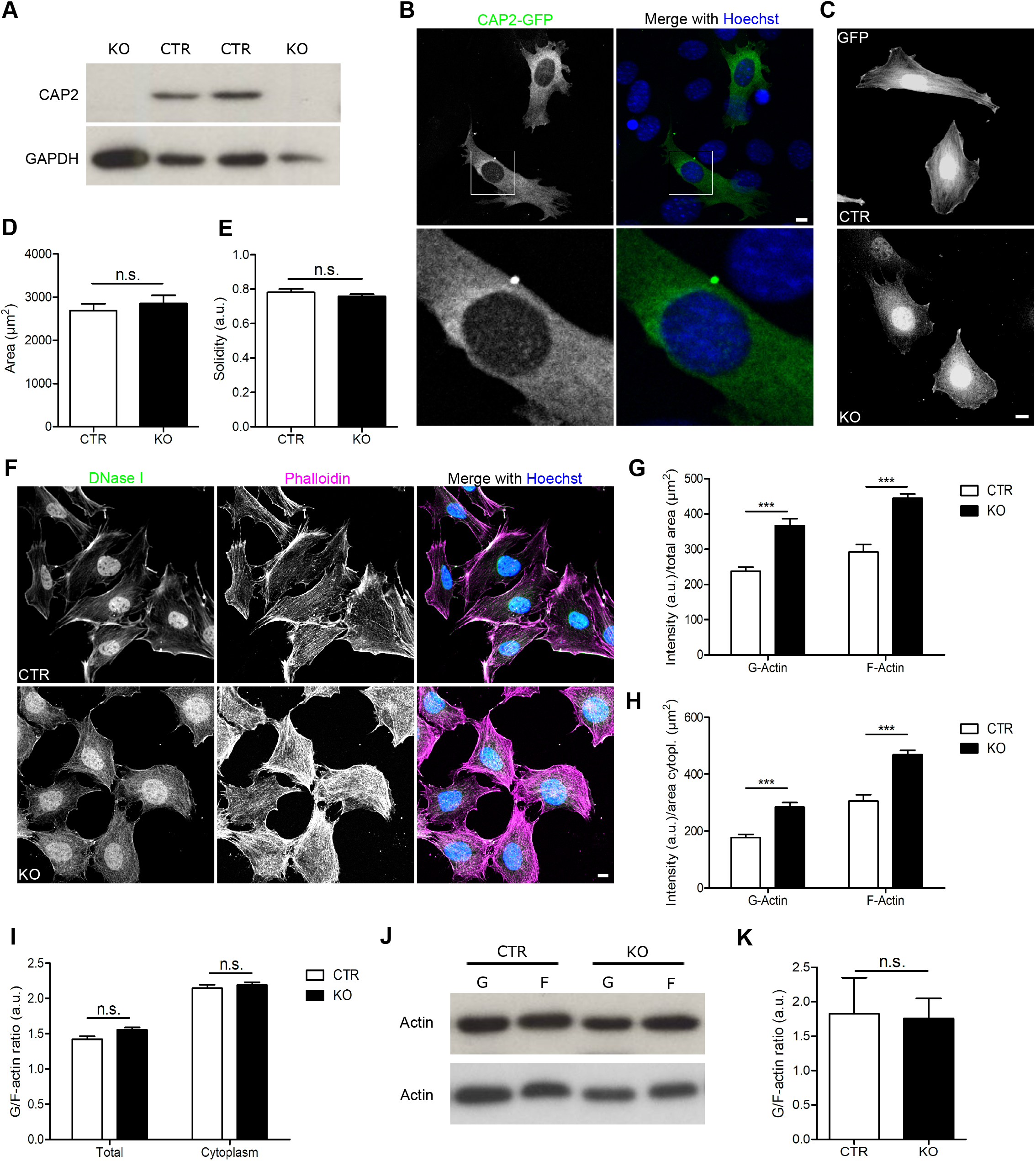
CAP2 inactivation increased G-actin levels in MEFs. **(A)** Immunoblots showing CAP2 expression in two CTR MEF lines and CAP2 inactivation in two KO MEF lines. GAPDH was used as loading control. **(B)** Representative micrograph showing localization of GFP-tagged CAP2 (green) in CTR MEFs. Merge micrograph includes counterstaining with DNA dye 4′,6-diamidino-2-phenylindole (DAPI, blue). Box indicates area shown at higher magnification. **(C)** Representative micrographs of GFP-transfected CTR and KO MEFs. **(D)** Area quantification of CTR and KO MEFs. **(E)** Solidity index of CTR and KO MEFs. **(F)** Representative micrographs of MEFs stained with fluorescent phalloidin and DNase I. **(G)** Whole cell fluorescence quantification for phalloidin and DNase I. **(H)** Quantification of phalloidin and DNase I fluorescence in cytosol. **(I)** Quantification of DNase I-to-phalloidin-ratio for whole cell and cytosolic fluorescence. **(J)** Immunoblots showing actin in soluble (G) and insoluble (F) protein fractions. **(K)** Quantification of G/F-actin ratio. Scale bars (in μm): 10 (B. C, F). *: P<0.05, **: P<0.01, ***: P<0.001, ns: not significant.

KO MEFs normally adhered to cell culture dishes and did not differ from CTR MEFs in size or solidity index that we calculated to assess cellular morphology (Fig. 1C-E: size (in μm^2^): CTR: 2,689.45±154.77, KO: 2,856.97±184.21, n≥25/3, P=0.500; solidity index (arbitrary units): CTR: 0.78±0.02, KO: 0.76±0.01, n≥25/3 cells, P=0.316). Further, real-time impedance measurements revealed no differences between CTR and KO MEFs in cell proliferation (Fig. S1A), and CAP2 inactivation did not affect metabolic activity, mitochondrial morphology, mitochondrial ROS production (Fig. S1E), lipid peroxidation or mitochondrial membrane potential (Fig. S1B-G). Furthermore, oxidative stress induced by erastin treatment similarly changed metabolic and mitochondrial parameters in CTR and KO MEFs (Landshamer et al. 2008; Neitemeier et al. 2017). Together, morphology, proliferation, metabolic activity, mitochondrial function and response to oxidative stress were unchanged in KO MEFs. Hence, CAP2 inactivation did not induce any obvious cellular defects.

Next, we studied whether CAP2 inactivation affected the actin cytoskeleton. To do so, we exploited fluorescent phalloidin that specifically labels F-actin and fluorescent DNase I that binds G-actin with high affinity (Cramer et al. 2002; Melak et al. 2017). In line with overall normal cellular functions, KO MEFs did not display any severe actin cytoskeleton defects (Fig. 1F). However, compared to CTR MEFs, fluorescent intensities of phalloidin and DNase I were both moderately increased in KO MEFs (Fig. 1G; DNase I: CTR: 237.41±11.62, KO: 366.63±19.83, n≥71/3, P<0.001; phalloidin: CTR: 292.02±21.41, KO: 444.60±12.09, n≥71/3, P<0.001). In CTR MEFs as well as in KO MEFs, DNase I fluorescence was much higher in the nucleus when compared to the cytosol and this likely did not reflect subcellular G-actin distribution.

We therefore left aside the nucleus and determined phalloidin and DNase I fluorescence solely in the cytosol. Similar to our analysis of total MEF fluorescent intensity, cytosolic phalloidin and DNase I fluorescence intensity was both higher in KO MEFs when compared to CTR MEFs (Fig. 1H; DNase I: CTR: 177.11±10.61, KO: 284.49±15.75, n≥71/3, P<0.001; phalloidin: CTR: 305.65±21.91, KO: 468.13±15.60, n≥71/3, P<0.001). However, the G/F-actin ratio calculated from DNase I and phalloidin fluorescence intensities was not different between CTR and KO MEFs independent of considering fluorescence intensity in the cytosol or in the entire cells (Fig. 1I; total: CTR: 1.42±0.04, KO: 1.56±0.03, n≥71/3, P=0.497; cytosol: CTR: 2.15±0.05, KO: 2.19±0.04, n≥71/3, P=0.880). A normal G/F-actin in KO MEFs was confirmed by immunoblots, for which we separated soluble protein from insoluble protein fractions including G-actin and F-actin, respectively (Fig. 1J-K; CTR: 1.83±0.47, KO: 1.76±0.26, n=5, P=0.916). Together, CAP2 inactivation did not induce severe defects in the actin cytoskeleton. However, it caused a moderate increase in G-actin and F-actin levels, which did not change the equilibrium between G-actin and F-actin.

### CAP2 inactivation alters subcellular distribution of MRTF-A in mouse fibroblasts

MRTF-A is a transcriptional coactivator that shuttles between the cytosol and the nucleus. This shuttling depends on G-actin, because G-actin binding is necessary for nuclear MRTF-A export and interferes with accessibility of its nuclear localization sequence (Plessner and Grosse 2015; Treisman 2013). Hence, moderately increased G-actin levels in KO MEFs might be associated with altered subcellular MRTF-A distribution. To test this, we generated CTR and KO MEF lines that stably expressed GFP-tagged MRTF-A (MRTF-A-GFP) and grouped MEFs into three categories: 1) MEFs with mainly nuclear MRTF-A-GFP (nuclear), 2) MEFs with mainly cytosolic MRTF-A-GFP (cytosolic) and 3) MEFs with equal MRTF-A-GFP levels in both compartments (equal), similar to previous studies (Hinojosa et al. 2017). In KO MEFs, MRTF-A-GFP distribution was different from CTR MEFs, as indicated by an almost sevenfold increase in the cytosolic MRTF-A-GFP fraction and a sevenfold decrease in the nuclear MRTF-A-GFP fraction (Fig. 2A, C; (in %) CTR: nuclear: 68.04±3.74, cytosolic: 11.51±2.58, equal: 20.45±2.22, n=15/3, KO: nuclear: 9.00±2.09, cytosolic: 85.22±2.61, equal: 5.78±1.95, n=15/3, P<0.001).

**Figure 2.**
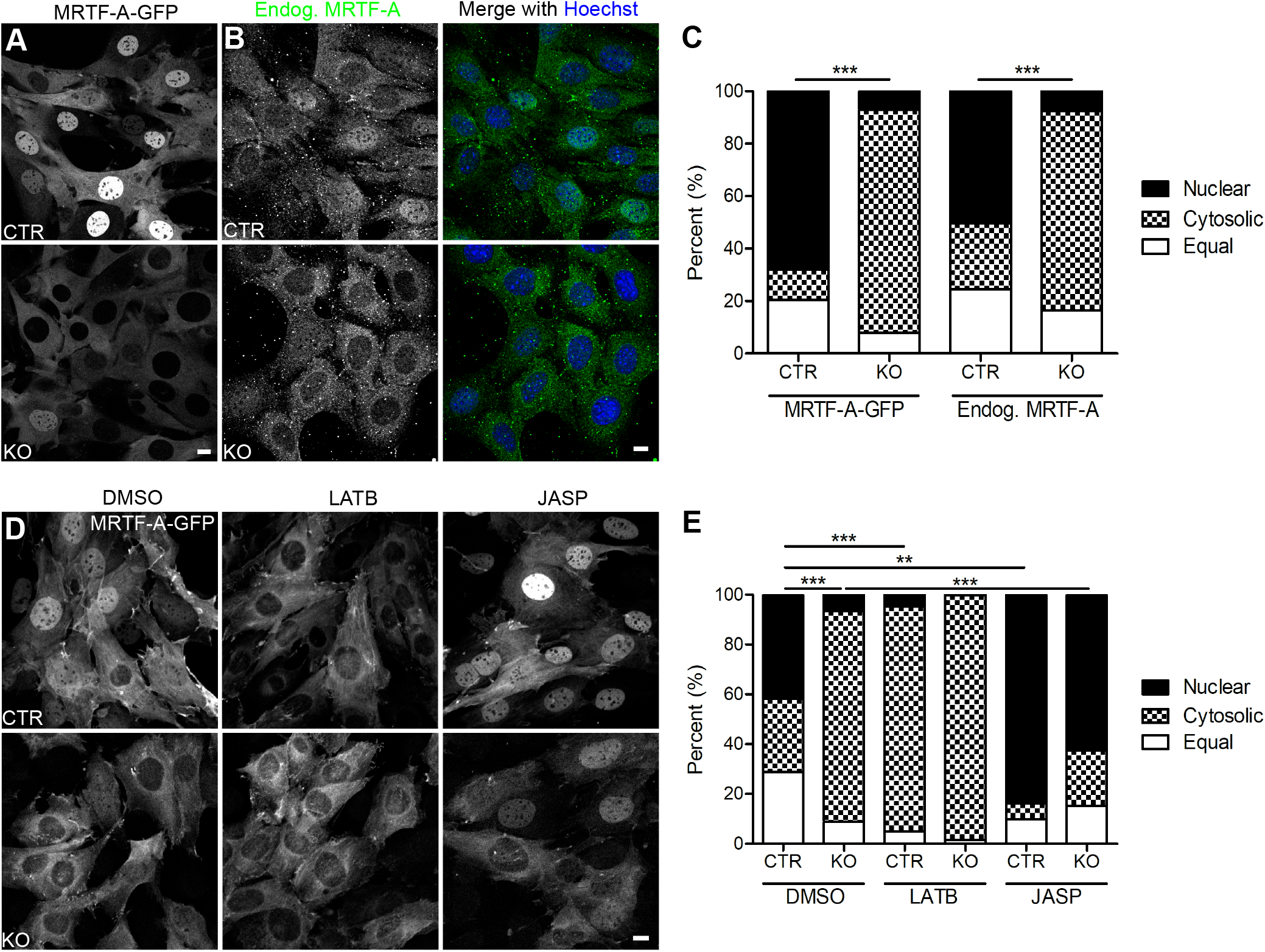
CAP2 controls MRTF-A localization in an actin-dependent manner. **(A)** Representative micrographs of CTR and KO MEFs that stably expressed GFP-tagged MRTF-A (MRTF-A-GFP). **(B)** Representative micrographs of CTR and KO MEFs stained with an antibody against MRTF-A (green). MEFs were counterstained with the DNA-dye Hoechst (blue). **(C)** Categorization of MEFs according to the localization of MRTF-A-GFP or endogenous MRTF-A, i.e. fractions with mainly nuclear or cytosolic MRTF-A-GFP and fraction with equal levels in both compartments. **(D)** Representative micrographs of MRTF-A-GFP-expressing CTR and KO MEFs upon treatment with either DMSO, latrunculin B (LATB) or jasplakinolide (JASP). **(E)** Categorization of MEFs according to the localization of MRTF-A-GFP upon treatment with either DMSO, LATB or JASP. Scale bars (in μm): 10 (A, B, D). *: P<0.05, **: P<0.01, ***: P<0.001, ns: not significant.

In an independent experiment, we determined localization of endogenous MRTF-A by immunocytochemistry and found very similar differences between CTR and KO MEFs (Fig. 2B-C; (in %) CTR: nuclear: 50.24±6.87, cytosolic: 25.23±3.60, equal: 24.53±4.77, n=6/3, KO: nuclear: 7.50±2.12, cytosolic: 75.97±4.38, equal: 16.53±3.24, n=6/3, P<0.001), thereby demonstrating that our GFP-tagged construct faithfully reflected MRTF-A localization and that our stably transfected MEF lines were valuable tools to study the mechanisms that control MRTF-A localization. Together, increased actin levels in KO MEFs were associated with altered MRTF-A localization, and we therefore hypothesized that CAP2 controls MRTF-A localization in an actin-dependent manner.

### F-actin stabilization restored MRTF-A localization in CAP2-deficient MEFs

To test this hypothesis, we exploited MRTF-A-GFP expressing MEF lines to determine MRTF-A localization upon pharmacological manipulation of the actin cytoskeleton. Latrunculin B (LATB) reportedly increased G-actin levels and promoted interaction of G-actin with MRTF-A (Morton et al. 2000). Hence, LATB treatment should increase cytosolic MRTF-A localization (Olson and Nordheim 2010). Indeed, when compared to dimethyl sulfoxide (DMSO)-treated CTR MEFs, LATB increased the CTR MEF fraction with cytosolic MRTF-A-GFP localization and it substantially decreased the fraction with nuclear localization (Fig. 2D-E; DMSO: nuclear: 41.76±2.68, cytosolic: 29.40±2.64, equal: 28.85±2.25, LATB: nuclear: 4.95±2.36, cytosolic: 90.20±3.43, equal: 4.85±1.86, n=9/3, P<0.01). In contrast, LATB failed in changing the subcellular MRTF-A-GFP distribution in KO MEFs (DMSO: nuclear: 6.74±1.43, cytosolic: 84.46±2.77, equal: 8.81±1.87, LATB: nuclear: 0.28±0.26, cytosolic: 98.35±0.64, equal: 1.38±0.66, n=9/3, P=0.158). Notably, subcellular MRTF-A distribution was not different between CTR and KO MEFs upon LATB treatment (P=0.225). Hence, LATB caused a subcellular MRTF-A distribution in CTR MEFs that was similar to that in KO MEFs.

Apart from LATB, we tested jasplakinolide (JASP) that stabilizes F-actin, reduces G-actin levels and that reportedly induced nuclear import of MRTF-A (Olson and Nordheim 2010; Tsuji et al. 2009). As expected, JASP doubled the fraction of CTR MEFs with nuclear MRTF-A-GFP localization and reduced the fraction with cytosolic localization (Fig. 2D-E; JASP: nuclear: 83.69±2.61, cytosolic: 6.41±1.05, equal: 9.90±2.16, n=9/3, P<0.01).

Similarly, JASP changed the subcellular MRTF-A-GFP distribution in KO MEF as indicated by a ninefold increased fraction with nuclear MRTF-A-GFP localization concomitant with a fourfold decreased cytosolic fraction (JASP: nuclear: 62.21±6.32, cytosolic: 22.28±5.95, equal: 15.50±2.19, n=9/3, P<0.01). Notably, the subcellular MRTF-A distribution did not differ between CTR and KO MEFs upon JASP treatment (P=0.237). Hence, pharmacologically induced reduction of G-actin levels restored MRTF-A localization in KO MEFs. These data suggested that CAP2 controlled MRTF-A localization in an actin-dependent manner.

### CAP2 is dispensable for serum induced nuclear MRTF-A translocation

Serum stimulation reportedly induced nuclear MRTF-A translocation in fibroblasts (Esnault et al. 2014; McGee et al. 2011; Miralles et al. 2003). By exploiting MRTF-A-GFP expressing MEF lines, we next tested whether CAP2 was relevant for serum-induced nuclear import of MRTF-A. First, we starved MEFs in 0.3% fetal calf serum (FCS) for 48 h and, thereafter, determined nuclear translocation during the first six min of stimulation with 20% FCS by live-cell imaging. We restricted this analysis to CTR and KO MEFs of the ‘cytosolic MRTF-A fraction’ to avoid any false interpretation due to different MRTF-A localization before stimulation. As obvious from the movies and image sequences (Fig. 3A, Movies S1-2), MRTF-A rapidly translocated into the nucleus in CTR and KO MEFs upon serum stimulation. Quantification of the latency of nuclear translocation revealed no difference between both groups (Fig 3B; (in s) CTR: 248.75±31.25, KO: 185.00±14.18, n=12/3, P=0.089). Next, we determined subcellular MRTF-A localization in all CTR and KO MEFs both upon 48 h of serum starvation and upon 10 min of serum stimulation. Compared to basal conditions (Fig. 2C), the CTR MEF fraction with nuclear MRTF-A localization was reduced by 40%, and the fraction with cytosolic MRTF-A was increased threefold upon starvation (Fig. 3C-D; nuclear: 40.90±3.78, cytosolic: 34.73±5.02, equal: 24.38±3.63, n=25/3, P<0.001). As expected, serum stimulation induced a nuclear translocation of all MRTF-A in CTR MEFs, and we did not note CTR MFEs with predominantly cytosolic MRTF-A or equal localization in cytosol and nucleus (Fig. 3C-D; n=9/3, P<0.001). In contrast to CTR MFEs, serum starvation did not alter MRTF-A localization in KO MEFs (nuclear: 1.67±1.07, cytosolic: 92.34±2.39, equal: 6.00±2.29, n=15/3, P<0.0543), while serum stimulation induced nuclear MRTF-A translocation in the majority of KO MEFs.

**Figure 3.**
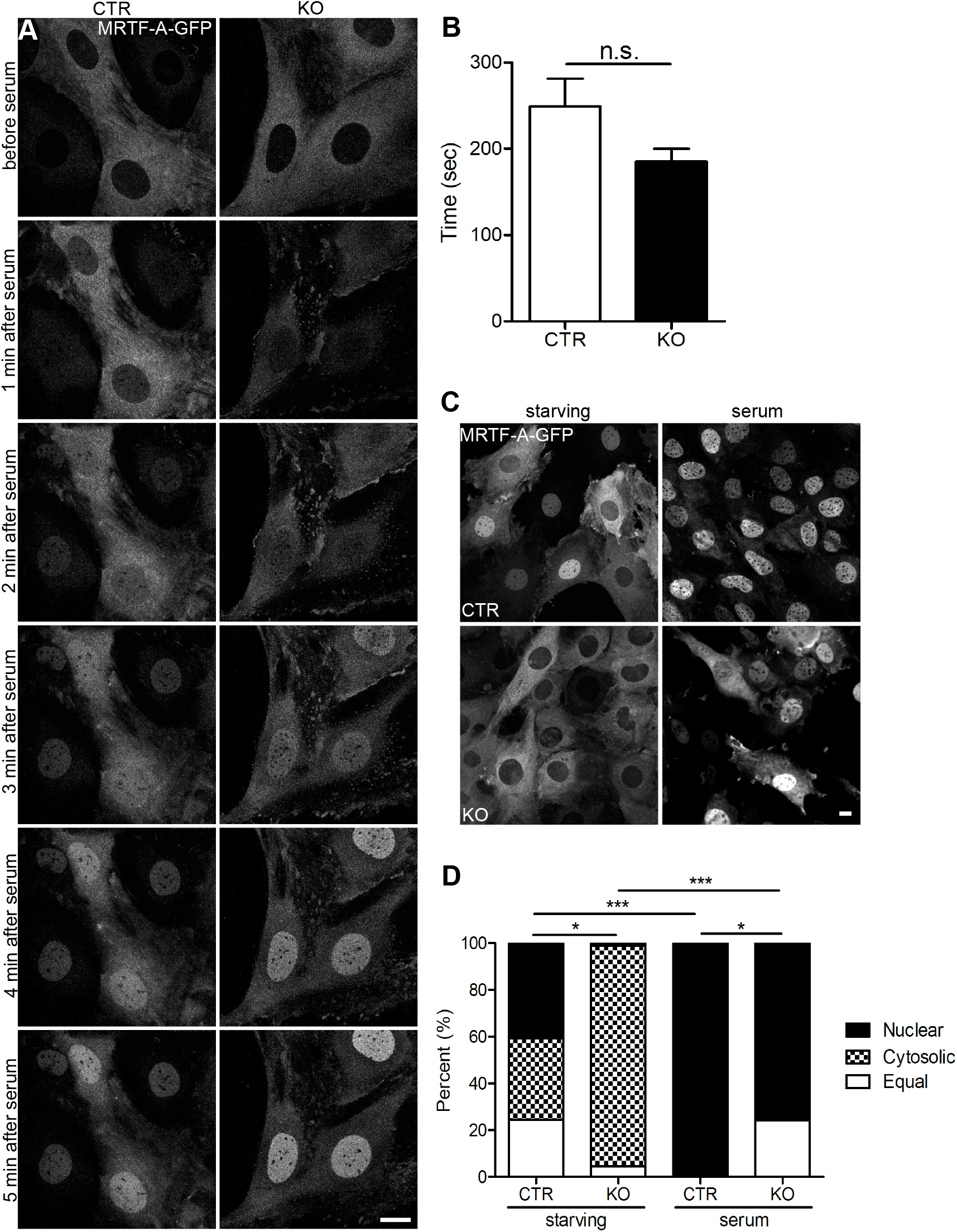
CAP2 was dispensable for serum induced nuclear MRTF-A translocation in MEFs. **(A)** Image sequence of MRTF-A-GFP-expressing MEFs before and during serum stimulation. **(B)** Latency of nuclear MRTF-A translocation. **(C)** Representative micrographs of MRTF-A-GFP-expressing CTR and KO MEFs during serum starvation and upon serum stimulation. **(D)** Categorization of MEFs according to the localization of MRTF-A-GFP or endogenous MRTF-A during serum starvation and upon serum stimulation. Scale bars (in μm): 10 (A, C). *: P<0.05, **: P<0.01, ***: P<0.001, ns: not significant.

However, unlike in FCS-stimulated CTR MEFs, we noted a fraction of 25% KO MEFs with equal localization of MRTF-A in cytosol and nucleus upon FCS stimulation (nuclear: 75.02±3.79, equal: 24.98±3.79, n=9/3, P<0.001). Upon serum stimulation, subcellular MRTF-A distribution was still different between CTR and KO MEFs (P<0.05). Together, serum stimulation induced nuclear MRTF-A translocation in KO MEFs with a latency similar to CTR MEFs. However, different from CTR MEFs, in a quarter of serum-stimulated KO MEFs MRTF-A was still present in the cytosol.

### CAP2 inactivation reduces SRF activity in mouse fibroblasts

So far, we showed that CAP2 i) controls MRFT-A localization in MEFs and ii) was largely dispensable for serum-induced nuclear MRTF-A translocation. Next, we tested whether CAP2 was relevant for MRFT-A-dependent activation of SRF. To do so, we generated CTR and KO MEF lines that stably expressed a SRF reporter in which expression of firefly luciferase was controlled by three minimal c-Fos promotor sequences including serum response elements, but lacking TCF binding sites (Hinojosa et al. 2017). Luciferase activity was strongly reduced in KO MEFs and reached only half of that in CTR MEFs, thereby suggesting reduced SRF activity in KO MEFs (Fig. 4A; (arbitrary units) CTR: 48.19±6.94, KO: 27.28±1.81, n=6, P<0.05). Indeed, qPCR experiments revealed a reduction in mRNA levels for several established SRF downstream target genes including those encoding for c-Fos (*c-Fos*), vinculin (*Vcl*), smooth muscle α-actin (*Acta2*), Cyr61 (*Cyr61*) or SRF (*Srf*) itself (Fig. 4B). In these experiments, we found similar changes in both KO MEF lines generated (KO 1: *Cyr61*: 0.16±0.02, P<0.01; *c-Fos*: 0.16±0.03, P<0.05; *Acta2*: 0.29±0.05, P<0.05; *Vcl*: 0.20±0.05, P<0.05; *Srf*: 0.48±0.07, P=0.073, n=9; KO 2: *Cyr61*: 0.48±0.11, P<0.05; *Fos*: 0.11±0.02, P<0.05; *Acta2*: 0.11±0.02, P<0.05; *Vcl*: 0.24±0.06, P<0.05; *Srf*: 0.29±0.04, P<0.05, n=9). Notably, mRNA levels of *Egr2*, a SRF downstream target that is controlled by the TCF, but not by the MRTF-A pathway, was unchanged in both KO MEF lines (KO 1: 1.03±0.20, n=9, P=0.923; KO 2: 0.70±0.03, n=9, P=0.319). Together, altered MRTF-A localization in KO MEFs was associated with reduced SRF activity.

**Figure 4.**
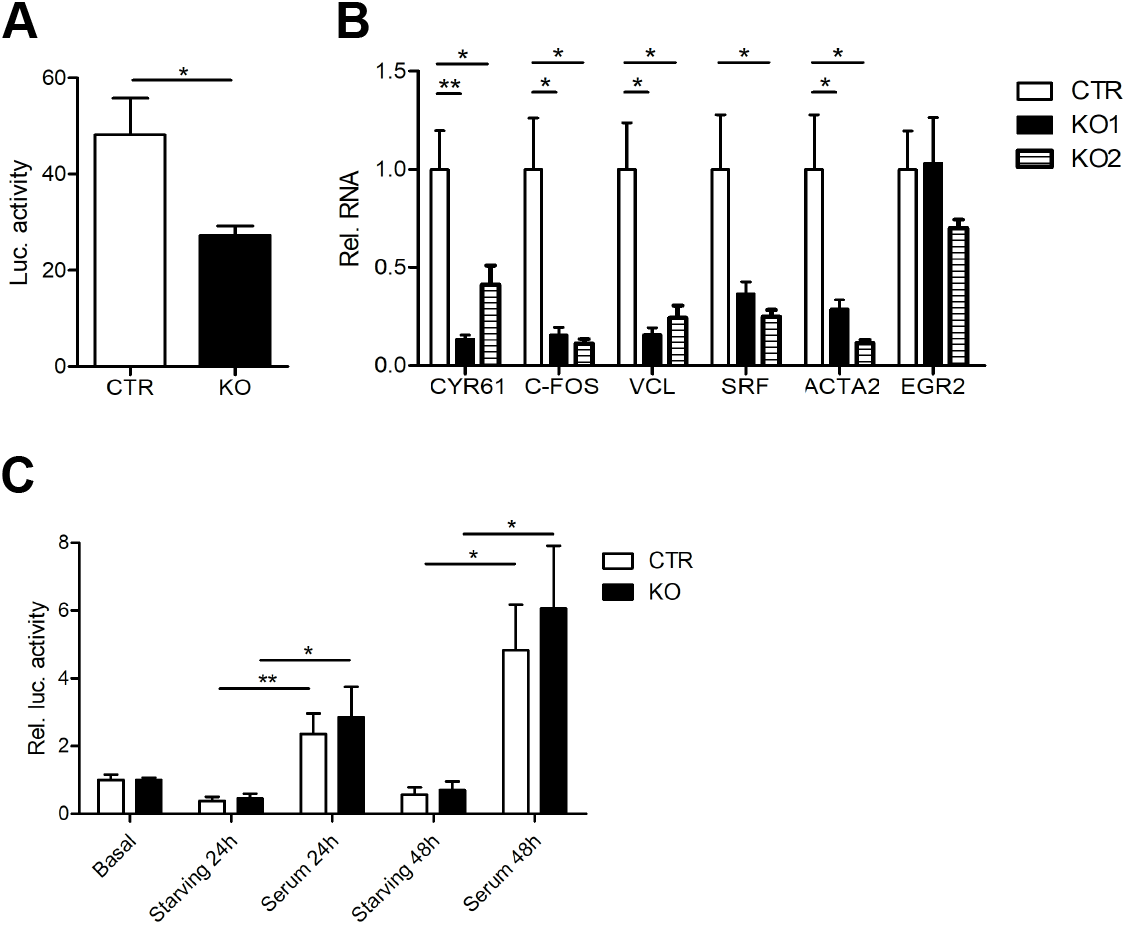
CAP2 inactivation reduced SRF activity in MEFs. **(A)** Firefly luciferase activity in CTR and KO MEFs that stably expressed a SRF reporter in which firefly luciferase expression was under control of SRF activity. **(B)** mRNA levels of selected SRF target genes in two KO MEF lines as determined by qPCR. **(C)** Luciferase expression in CTR and KO MEFs under basal conditions, during starving and upon serum stimulation for either 24 or 48 h. Values are normalized to basal levels. *: P<0.05, **: P<0.01, ***: P<0.001, ns: not significant.

Next, we wanted to know whether serum-induced SRF activity depends on CAP2. To test this, we starved CTR and KO MEFs expressing the SRF reporter for 20 h before stimulation with 20% FCS for either 24 or 48 h.

As expected, serum stimulation for either 24 or 48 h induced SRF activity in CTR MEFs (Fig. 4C; 24 h: 2.36±0.55, n=6, P<0.01; 48 h: 4.83±1.22, n=6, P<0.05). Very similar to CTR MEFs, serum stimulation induced SRF activity in KO MEFs (24 h: 2.86±0.81, n=6, P<0.05; 48 h: 6.06±1.69, n=6, P<0.05), and serum induced SRF activity was not different between groups (24 h: P=0.652; 48 h: P=0.601). Together, basal SRF activity was reduced in KO MEFs, but serum stimulation equally induced SRF activity in CTR and KO MEFs.

## Discussion

The present study aimed at deciphering the CAP2-dependent mechanism relevant for the control of the transcription factor SRF. We chose mouse embryonic fibroblasts (MEFs) as a cellular model system for our study, because in these cells CAP2 is expressed at substantial levels and SRF-dependent gene regulation and upstream regulatory mechanisms have been intensively studied (Esnault et al. 2014). We found increased DNase I fluorescent intensity in CAP2 mutant MEFs, which was associated with a reduction in nuclear MRTF-A, reduced SRF activity and decreased expression of established MRTF-SRF target genes. While drug-induced increase in G-actin levels altered MRTF-A localization in control MEFs, it did not affect MRTF-A localization in mutant MEFs, which was normalized upon drug-induced decrease in G-actin levels. These data let us propose a model in which CAP2 controls SRF-dependent gene expression via regulating G-actin levels and nuclear MRTF-A localization.

This model is in good agreement with recent studies that identified important functions for CAPs in controlling F-actin dynamics (Chaudhry et al. 2010; Jansen et al. 2014; Johnston et al. 2015; Kotila et al. 2018; Kotila et al. 2019; Mu et al. 2019; Shekhar et al. 2019). Specifically, these studies showed that CAPs can i) facilitate F-actin severing and actin subunit dissociation in synergy with ADF/cofilin and twinfilin (Chaudhry et al. 2010; Jansen et al. 2014; Johnston et al. 2015; Kotila et al. 2018; Kotila et al. 2019; Shekhar et al. 2019), ii) catalyze ATP-for-ADP exchange on G-actin that is relevant for F-actin assembly (Chaudhry et al. 2007; Jansen et al. 2014; Moriyama and Yahara 2002; Nomura et al. 2012), and iii) inhibit activity of inverted formin 2 (INF2) that promotes F-actin assembly (Mu et al. 2019). Hence, by regulating various aspects of F-actin assembly and disassembly, CAPs control G-actin levels and, hence, interaction of G-actin with MRTF-A. Our finding of increased DNase I fluorescent intensity and reduced nuclear MRTF-A localization in CAP2 mutant MEFs, together with normalization of MRTF-A localization upon treatment with the F-actin stabilizing drug JASP emphasize that CAP2 inactivation increased G-actin levels in MEFs, suggesting that in MEFs CAP2 acts as a F-actin assembly factor rather than promoting F-actin disassembly.

We found reduced SRF activity in reporter assays as well as reduced expression of SRF target genes in CAP2 mutant MEFs.

Supportively, reduced expression of MRTF-SRF targets in CAP2 mutant MEFs have been noted recently by others (Xiong et al. 2019). Although SRF activity has not been systematically analyzed in skeletal muscle from CAP2 mutant mice, decreased mRNA levels of established MRTF-SRF targets such as *Acta1*, *Acta2* and *Actc1* point towards reduced SRF activity in skeletal muscles from CAP2 mutant embryos, too (Kepser et al. 2019), in which SRF dysregulation may contribute to retarded postnatal skeletal muscle development. Indeed, a crucial function for the MRTF-SRF pathway during late embryonic skeletal muscle development is evident from gene-targeted mice (Cenik et al. 2016; Li et al. 2005). Opposite to our findings in CAP2 mutant MEFs and to reduced expression of MRTF-SRF target genes in skeletal muscles from CAP2 mutant embryos (Kepser et al. 2019), a recent study reported upregulation of several MRTF-SRF target genes including *Acta1* and *Acta2* in heart tissue and isolated cardiomyocytes from CAP2 mutant mice, which was associated with increased nuclear MRTF levels. Interestingly, heart defects in CAP2 mutant mice including dilated cardiomyopathy and impaired cardiac conductance were partially restored upon pharmacological inhibition of MRTF-SRF activity, demonstrating that CAP2-dependent regulation of the MRTF-SRF pathway is physiologically relevant and that its dysregulation due to CAP2 inactivation contributes to pathological conditions (Xiong et al. 2019). The opposite effects of CAP2 inactivation on MRTF-SRF activity suggest different CAP2 activities towards F-actin dynamics in MEFs versus cardiomyocytes. Our data suggest that CAP2 promotes F-actin assembly in MEFs. Instead, it may primarily act as a F-actin disassembly factor in cardiomyocytes.

By chromatin immunoprecipitation combined with deep sequencing, previous studies convincingly demonstrated that serum-induced, SRF-mediated transcriptional response largely depends on the MRTF-SRF pathway (Esnault et al. 2014). In line with these data, we showed efficient MRTF-A translocation into the nucleus as well as elevated SRF activity upon serum stimulation in control MEFs. Interestingly, serum-stimulated nuclear MRTF-A translocation as well as SRF activation was similar to controls in CAP2 mutant MEFs. Hence, while we found a role for CAP2 in MRTF-A localization and SRF activity under basal conditions, in unstimulated MEFs, CAP2 was dispensable for serum stimulation of the MRTF-SRF pathway.

Via acting on transmembrane receptors including G-protein-coupled receptors, receptor tyrosine kinases or serine-threonine receptor kinases, extracellular signals are translated into intracellular signaling cascades that include Rho family small guanosine triphosphatases (GTPases) (Cotton and Claing 2009; Moustakas and Heldin 2008; Schiller 2006). Effectors of Rho GTPases include, among others, formins, Wiskott-Aldrich syndrome protein (WASP), WASP-family verprolin homologues (WAVEs) and actin-related protein 2/3 (ARP2/3) complex, which orchestrate actin polymerization (Jaffe and Hall 2005). Hence, Rho GTPase signaling promotes incorporation of G-actin into filaments that releases MRTF from G-actin complexes and stimulates MRTF-SRF-dependent gene expression (Olson and Nordheim 2010). Rho GTPase signaling further shifts the F/G-actin equilibrium towards F-actin via activation of Rho-associated kinases (ROCKs) that in turn inhibits actin depolymerizing proteins of the ADF/cofilin family (Ohashi et al. 2000). ADF/cofilin cooperates with CAPs in actin dynamics (Chaudhry et al. 2010; Jansen et al. 2014; Kotila et al. 2018; Kotila et al. 2019; Ono 2013), and inhibition of ADF/cofilin activity upon serum simulation may explain why CAP2 was dispensable for serum-induced stimulation of the MRTF-SRF pathway.

In summary, our data revealed that CAP2 controls the subcellular localization of MRTF and thereby SRF activity in unstimulated MEFs, while CAP2 was dispensable for serum-induced nuclear MRTF translocation and MRTF-SRF stimulation.

## Acknowledgements

We thank Maria Bettina Kowalski, Denis Grabski and Andrea Wüstenhagen for excellent technical support. This work was supported by a research grant from Fondazione Cariplo (2018-0511). LJK and LSH were supported by the DFG Research Training Group ‘Membrane Plasticity in Tissue Development and Remodeling’ (GRK 2213).

## Author contributions

Experiments were designed and results were discussed by LJK, LSH, CM, MR, EM, CC, RG and MBR. Data were analyzed by LJK, LSH, and CM. Manuscript was written by MBR and LJK.

---

**Figure S1.**
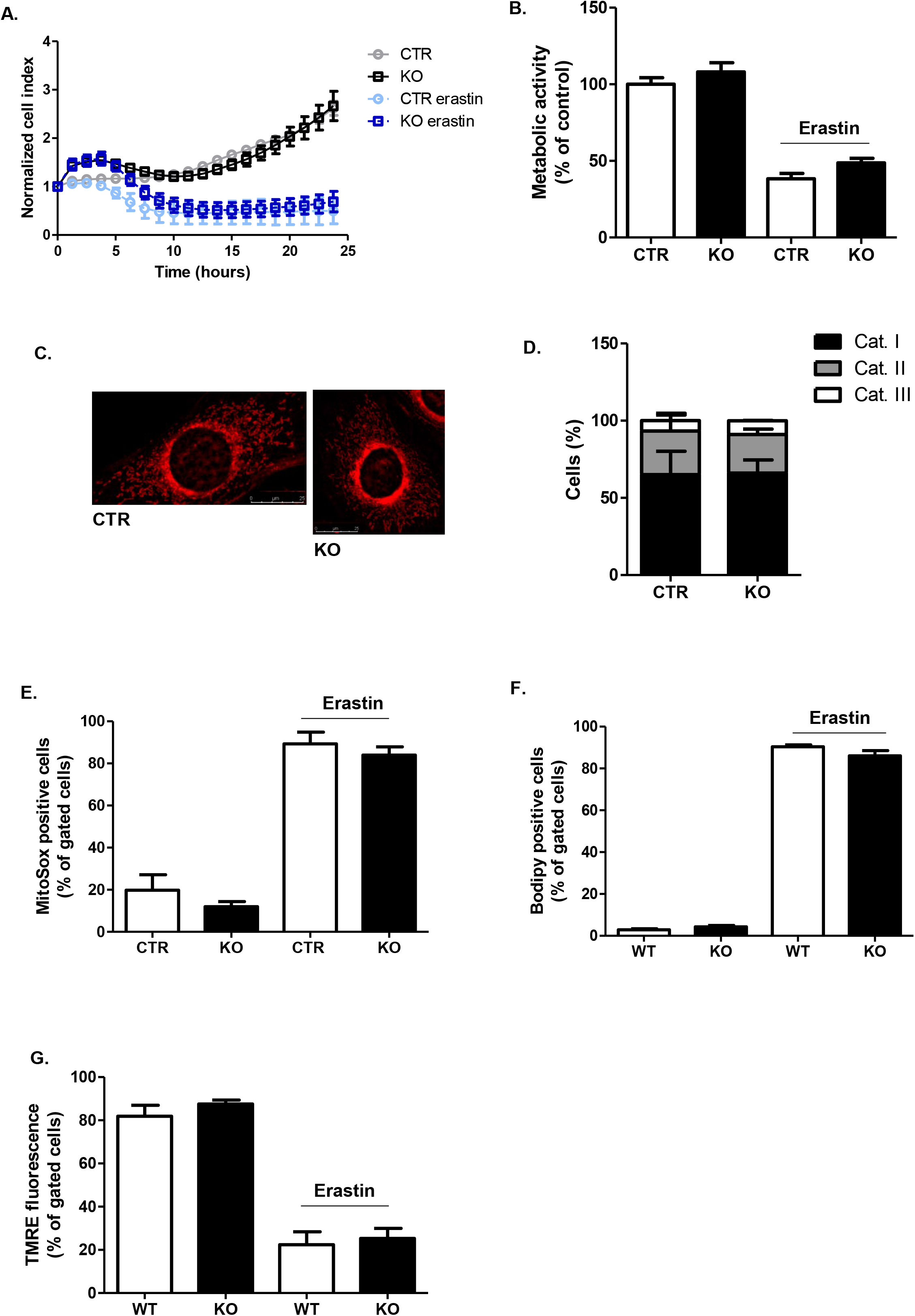
CAP2 inactivation does not affect energy metabolism and mitochondrial resilience in MEFs. **(A)** Real-time cell impedance measurements performed in CTR and KO MEFs. Cells were treated or not with erastin (0.7μM). **(B)** Metabolic activity was determined by MTT assay. CTR and KO MEFs were challenged or not with erastin (0.7μM) for eight hours and were presented as percentage of CTR. **(C)** Representative micrographs of CTR and KO MEFs stained with MitoTracker Deep Red to visualize mitochondrial morphology. Scale bars: 25μm. **(D)** Analysis of mitochondrial fragmentation performed in five hundred CTR and KO MEF cells quantified as percentage of counted cells. **(E)** MitoSOX staining and subsequent FACS analysis were performed in CTR and KO MEFs treated or not with erastin (0.7μM) for eight hours. **(F)** BODIPY 581/591 staining and subsequent FACS analysis were performed in CTR and KO MEFs treated or not with erastin (0.7μM) for eight hours. **(G)** TMRE staining and subsequent FACS analysis were performed in CTR and KO MEFs treated or not with erastin (0.7μM) for sixteen hours. For panels E, F and G, data are presented as percentage of gated cells and are representative of three replicates per condition.

